# Genomic Resources for Asian (*Elephas maximus*) and African Savannah Elephant (*Loxodonta africana*) Conservation and Health Research

**DOI:** 10.1101/2023.02.10.528067

**Authors:** Natalia A. Prado, Ellie E. Armstrong, Janine L. Brown, Shifra Z. Goldenberg, Peter Leimgruber, Virginia R. Pearson, Jesús E. Maldonado, Michael G. Campana

## Abstract

We provide novel genomic resources to help understand the genomic traits involved in elephant health and to aid conservation efforts. We sequence 11 elephant genomes (5 African savannah, 6 Asian) from North American zoos, including 9 *de novo* assemblies. We estimate elephant germline mutation rates and reconstruct demographic histories. Finally, we provide an in-solution capture assay to genotype Asian elephants. This assay is suitable for analyzing degraded museum and non-invasive samples, such as feces and hair. The elephant genomic resources we present here should allow for more detailed and uniform studies in the future to aid elephant conservation efforts and disease research.

## Introduction

Elephants are facing extinction. Approximately 50,000 Asian elephants (*Elephas maximus*) remain in fragmented populations across 13 range states in Asia (Williams et al. 2020). African elephants comprise two species (Roca et al. 2001; Rohland et al. 2010): African savannah elephants (*Loxodonta africana*) and African forest elephants (*Loxodonta cyclotis*). Population numbers of African elephants have dropped precipitously since the early 2000s: savannah and forest elephant populations have declined 30% and 62% respectively to a combined total of ~400,000 individuals (Chase et al. 2016; Gobush et al. 2021, 2022; Maisels et al. 2013; Thouless et al. 2016). Savannah elephants persist in 24 countries across Africa, inhabiting ~15% of their pre-agricultural range (Chase et al. 2016; Gobush et al. 2022). Fragmented forest elephant populations remain across 20 countries in the humid forest area of western and central Africa (Gobush et al. 2021). Asian and African savannah elephants are listed as Endangered by the International Union for the Conservation of Nature (IUCN) Red List of Threatened Species™ (Gobush et al. 2022; Williams et al. 2020), while African forest elephants are considered Critically Endangered (Gobush et al. 2021). Asian elephants are included in the Convention on International Trade in Endangered Species of Wild Fauna and Flora (CITES) Appendix I, while African elephants are listed under Appendices I or II depending on the population (CITES 2023).

Significant threats to elephant conservation include poaching and the loss and fragmentation of habitat due to human expansion, infrastructure development, and agricultural land conversion, which leads to human-elephant conflict (Gobush et al. 2021, 2022; Leimgruber et al. 2003, 2011; Williams et al. 2020). These threats may have downstream implications for population genetic structure, as range restriction may limit gene flow (Athira and Vidya 2021), and periods of heightened mortality or targeted killing directed at mature adults may lead to reconstituted social groups of lower relatedness or may limit the number of mature males contributing to the gene pool (Athira and Vidya 2021; Gobush et al. 2008; Wittemyer et al. 2009). In extreme cases, decreased genetic diversity may lead to inbreeding depression (Vidya et al. 2007). Further compounding conservation efforts are poor estimates of elephant population sizes and distribution ranges across Asian and African countries (Blake and Hedges 2004; Chase et al. 2016; Gobush et al. 2021, 2022; Maisels et al. 2013; Thouless et al. 2016; Williams et al. 2020). Access to technologies that can potentially address these estimates, such as genomics, is frequently limited in elephant-range countries (e.g. Helmy et al. 2016).

Captive breeding is increasingly viewed as a means of maintaining important populations as “insurance” against environmental or anthropomorphic catastrophe (Faust and Marti 2011a, 2011b; Hildebrandt et al. 2012; Lei et al. 2008, 2011; Olson and Weise 2000; Thitaram 2012). Unfortunately, the clear majority of captive elephant populations are not self-sustaining due to high mortality and low birth rates, and although occurring at a lower rate than in years past, importation of wild-caught or captive-born elephants continues to be a necessary tool in maintaining the genetic diversity of the population (Clubb and Mason 2002; Faust et al. 2011a, 2011b;

Hildebrandt et al. 2012; Olson and Weiss 2000; Prado-Oviedo et al. 2016; Thitaram 2012; Wiese 2000). Over 70% of the North American elephant captive population is wild-born, and less than 30% have produced offspring (Prado-Oviedo et al. 2016), raising concerns about the genetic health of future generations. Captive elephant sustainability has been thwarted by numerous reproductive and health problems, including ovarian cycle abnormalities, uterine pathologies, gestational difficulties, diseases, and bull subfertility (Brown 2014). Mitigation of these problems could benefit from increased knowledge of the genetic background of captive breeding populations and their wild counterparts in range countries (Lei et al. 2008, 2011).

Despite a substantial body of genetic and genomic research on elephants (e.g. Ahlering et al. 2012a, 2012b; Fleischer et al. 2001; Hauf et al. 2000; Lei et al. 2008, 2011; Palkopoulou et al. 2018; Roca et al. 2001; Rogaev et al. 2006; Rohland et al. 2010; Tollis et. al 2021), publicly available *de novo* elephant genome assemblies are limited. Using Sanger sequencing, the Broad Institute generated a chromosome-level African savannah elephant reference sequence (loxAfr4.0: Palkopoulou et al. 2018). Tollis et al. (2021) generated a *de novo* draft sequence of an Asian elephant (‘Icky’) using Illumina shortread mate-pair libraries of varying insert sizes. Recently, the DNA Zoo (https://www.dnazoo.org/assemblies/Elephas_maximus) and the Vertebrate Genomes Project (VGP; https://genomeark.github.io/genomeark-all/Elephas_maximus/) have released Asian elephant genome assemblies, using Hi-C and the VGP telomere-to-telomere assembly pipeline respectively (Aiden Lab 2018– 2022; VGP 2022). Neither of these assemblies has yet been published in a peer-reviewed journal, and the VGP assembly is subject to a strict data-use policy.

Here we describe novel genomic resources for elephant health and conservation through sequencing of genomes from 11 (6 Asian and 5 African savannah) elephants in the North American population. Nine of these elephant genomes (5 Asian and 4 African savannah) were assembled *de novo*, including a hybrid 10x and Hi-C reference assembly of an Asian elephant bull that is substantially more contiguous than the published assembly for ‘Icky’ (Tollis et al. 2021) in terms of scaffold (37×: 102.3 Mb versus 2.77 Mb for Icky) and contig N50 values (8.5×: 681.5 Kb versus 79.8 Kb for Icky). We resequenced another 2 elephants (1 Asian 1 African savannah), for a total of 11 genomes. Additionally, we estimated germline mutation rates for both species using parent-offspring trios and reconstructed demographic histories for all individuals. Finally, we developed an in-solution capture assay to genotype low-quality and non-invasive Asian elephant samples (e.g. fecal samples and museum specimens). This assay will enable conservation genomics efforts in areas where access to fresh, genome-quality tissues are limited.

## Methods

### Biological Materials

We analyzed 11 (6 Asian and 5 African savannah) elephants from six Association of Zoos and Aquariums (AZA)-accredited institutions in North America (Supplementary Table 1). We denote each individual using ‘Em’ for Asian elephants or ‘La’ for African savannah elephants, followed by the AZA elephant studbook number. Whole blood samples were collected from an ear or leg vein directly into EDTA tubes during regular veterinary examinations and shipped frozen to the Center for Conservation Genomics at the Smithsonian’s National Zoo and Conservation Biology Institute (NZCBI). The only exception was the Asian elephant Em147 sample, which was obtained from the biobank at the National Elephant Herpes Laboratory. This sample had been collected for the 1998 artificial insemination of dam Em165 that resulted in the birth of Em538. All samples were stored in an ultralow (−80°C) freezer until DNA extraction.

### Nucleic acid library preparation

DNA was extracted from 250 μl aliquots of whole blood with the BioSprint 96 DNA Blood Kit (Qiagen Corp.) using the BioSprint 96 robot in 96-well format. DNA yield was measured using a Qubit high sensitivity, doublestranded DNA kit (Life Technologies), using 1 μL of input DNA. Library preparation using 10x Genomics Chromium chemistry was completed at HudsonAlpha Institute of Biotechnology (Huntsville, AL). For one elephant (Em538, “Kandula”), we built Dovetail Hi-C libraries (Cat #21004) according to the manufacturer’s instructions. For *de novo* mutation rate estimation using parent-offspring trios, two sire elephants, Em147 and La368, were resequenced at Fulgent Genetics (Temple City, CA) using single-indexed Illumina TruSeq libraries.

### DNA sequencing and genome assembly

We sequenced one elephant reference genome (Em538) to ~60× (2 lanes of Illumina Hiseq X Ten) coverage and eight additional female genomes (4 Asian and 4 African) at ~30× (1 lane of Illumina Hiseq X Ten) coverage using 10x Linked-Reads at HudsonAlpha Institute of Biotechnology. The Dovetail library for Em538 was sequenced at Admera Health (South Plainfield, NJ) on one lane of Illumina Hiseq X Ten. The two resequenced sire elephants were 2 × 150 paired-end sequenced at Fulgent Genetics to ~30× coverage on one lane of Illumina HiSeq X each.

For each of the nine 10x-sequenced individuals, we *de novo* assembled the genomes using Supernova 2.0.0 (Weisenfeld et al. 2017; See Table 1), specifying that all of the reads should be used (option ‘--maxreads=all’). Using the National Center for Biotechnology Information (NCBI) Foreign Contamination Screen (run August 2022: NCBI 2022) and the genome_decontaminate v. 0.4.0 script (Campana 2019–2022), we marked contigs corresponding to the mitogenome and removed duplicate contigs, adapter contaminations (splitting contigs where contaminations were identified), and sequences shorter than 200 bp. For Em538, we then scaffolded the 10x assembly using the Dovetail proximity ligation data using YaHS v. 1.1ar-3 (Zhao et al. 2023). Hereafter, we denote Em538’s 10x-only assembly as ‘10x’ and the Hi-C-scaffolded assembly as ‘10x+Hi-C’.

**Table 1.** Software used in the assembly and analysis of the *de novo* elephant genomes. Multiple versions of software are separated by a slash (/) and listed in the order they appear in the manuscript.

| Program | Version(s) | Citation |
| --- | --- | --- |
| Supernova | 2.0.0 | Weisenfeld et al. 2017 |
| NCBI Foreign Contamination Screen | August 2022 | NCBI 2022 |
| genome_decontaminate | 0.4.0 | Campana 2019–2022 |
| YaHS | 1.1a-r3 | Zhao et al. 2023 |
| assemblathon_stats.pl | 11 December 2022 | Bradnam et al. 2013 |
| BUSCO | 5.3.2 | Manni et al. 2021 |
| RepeatMasker | 4.0.9/4.0.7 | Smit et al. 2013–2015 |
| RMBlast | 2.9.0+ | Smit et al. 2013–2015 |
| RepeatModeler | 2.0.1 | Flynn et al. 2020 |
| GenMap | 1.2.0 | Pockrandt et al. 2020 |
| SeqAn | 2.4.1 | Reinert et al. 2017 |
| Exonerate | 2.2.0 | Slater and Birney 2005 |
| BLAT | 36 | Kent 2002 |
| AUGUSTUS | 3.3.2 | Stanke et al. 2006a, 2006b, 2008 |
| InterProScan | 5.60-92.0 | Jones et al. 2014 |
| RatesTools | 0.5.13 | Armstrong and Campana 2023 |
| Nextflow | 22.10.4.5836 | Di Tommaso et al. 2017 |
| Elephant Analysis Pipeline | 0.1.0 | This manuscript |
| BWA | 0.7.17-r1188 | Li 2013 |
| SAMtools | 1.13/1.6 | Danecek et al. 2021; Li et al. 2009 |
| Genome Analysis Toolkit | 4.3.0.0/4.0.8.1 | McKenna et al. 2010 |
| BCFtools | 1.13/1.6 | Danecek et al. 2021 |
| PSMC | 0.6.5 | Li and Durbin 2011 |
| Trimmomatic | 0.36 | Bolger et al. 2014 |
| Picard Tools | 2.5.0 | Broad Institute 2016 |
| VCFtools | 0.1.14/0.1.17 | Danecek et al. 2011 |
| BaitsTools | 1.6.6 | Campana 2018 |
| elephant_drop_baits.rb | 2020 January 31 | This manuscript |

### Assembly quality

We calculated genome assembly statistics for all assemblies (Tables 2–3; Supplementary Table 2) using the assemblathon_stats.pl script from the Assemblathon 2 competition (Bradnam et al. 2013). We assessed genic completeness using BUSCO v. 5.3.2 (options ‘-m geno --long -z’) with the mammalia_odb10 database (Manni et al. 2021; Table 4).

**Table 2.**
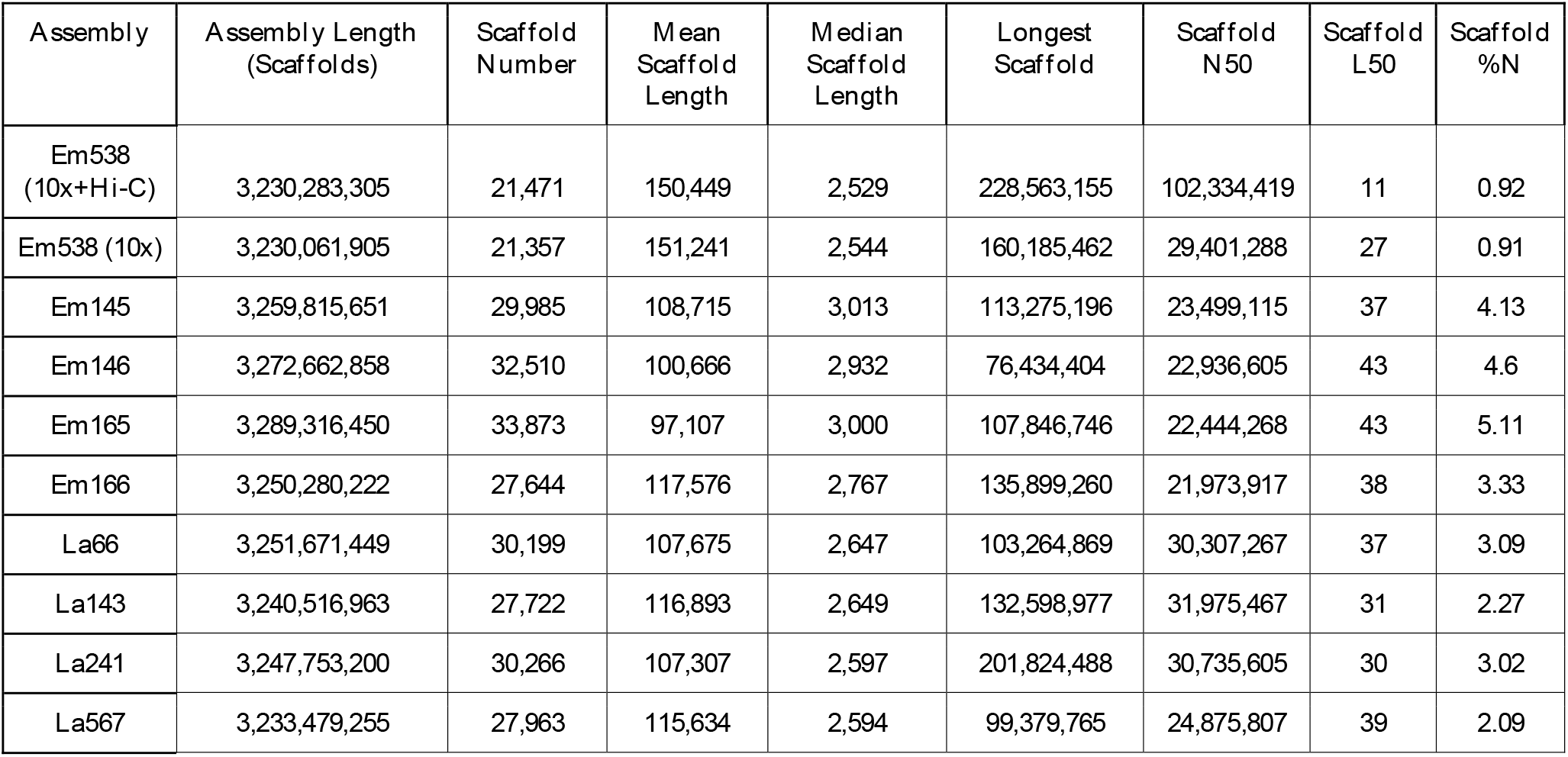
Scaffold-level genome assembly statistics for the *de novo* elephant assemblies. Statistics for our reference assembly (Em538) are given for both the 10x-only and the final 10x+Hi-C builds.

| Assembly | Assembly Length<br>(Scaffolds) | Scaffold<br>Number | Mean<br>Scaffold<br>Length | Median<br>Scaffold<br>Length | Longest<br>Scaffold | Scaffold<br>N50 | Scaffold<br>L50 | Scaffold<br>%N |
| --- | --- | --- | --- | --- | --- | --- | --- | --- |
| Em538<br>(10x+Hi-C) | 3,230,283,305 | 21,471 | 150,449 | 2,529 | 228,563,155 | 102,334,419 | 11 | 0.92 |
| Em538 (10x) | 3,230,061,905 | 21,357 | 151,241 | 2,544 | 160,185,462 | 29,401,288 | 27 | 0.91 |
| Em145 | 3,259,815,651 | 29,985 | 108,715 | 3,013 | 113,275,196 | 23,499,115 | 37 | 4.13 |
| Em146 | 3,272,662,858 | 32,510 | 100,666 | 2,932 | 76,434,404 | 22,936,605 | 43 | 4.6 |
| Em165 | 3,289,316,450 | 33,873 | 97,107 | 3,000 | 107,846,746 | 22,444,268 | 43 | 5.11 |
| Em166 | 3,250,280,222 | 27,644 | 117,576 | 2,767 | 135,899,260 | 21,973,917 | 38 | 3.33 |
| La66 | 3,251,671,449 | 30,199 | 107,675 | 2,647 | 103,264,869 | 30,307,267 | 37 | 3.09 |
| La143 | 3,240,516,963 | 27,722 | 116,893 | 2,649 | 132,598,977 | 31,975,467 | 31 | 2.27 |
| La241 | 3,247,753,200 | 30,266 | 107,307 | 2,597 | 201,824,488 | 30,735,605 | 30 | 3.02 |
| La567 | 3,233,479,255 | 27,963 | 115,634 | 2,594 | 99,379,765 | 24,875,807 | 39 | 2.09 |

**Table 3.** Contig-level genome assembly statistics for the *de novo* elephant assemblies. Statistics for our reference assembly (Em538) are given for both the 10x-only and the final 10x+Hi-C builds.

| Assembly | Assembly Length (Contigs) | Contig Number | Number of Scaffolded Contigs | Mean Contig Length | Median Contig Length | Longest Contig | Contig N50 | Contig L50 |
| --- | --- | --- | --- | --- | --- | --- | --- | --- |
| Em538 (10x+Hi-C) | 3,200,814,413 | 33,876 | 14,039 | 94,486 | 4,285 | 4,270,977 | 681,476 | 1,345 |
| Em538 (10x) | 3,200,814,413 | 32,658 | 13,034 | 98,010 | 4,062 | 4,270,977 | 691,955 | 1,331 |
| Em145 | 3,125,296,252 | 76,544 | 50,976 | 40,830 | 11,020 | 967,254 | 121,392 | 7,462 |
| Em146 | 3,122,339,184 | 84,312 | 56,979 | 37,033 | 10,677 | 884,336 | 106,509 | 8,492 |
| Em165 | 3,121,389,005 | 87,550 | 58,955 | 35,653 | 10,897 | 859,647 | 101,870 | 8,897 |
| Em166 | 3,142,282,515 | 67,456 | 43,778 | 46,583 | 10,348 | 1,162,655 | 146,207 | 6,216 |
| La66 | 3,151,305,234 | 63,072 | 36,797 | 49,964 | 7,078 | 1,394,877 | 187,703 | 4,942 |
| La143 | 3,167,093,748 | 55,289 | 31,306 | 57,283 | 5,742 | 1,693,190 | 239,687 | 3,894 |
| La241 | 3,149,751,953 | 62,458 | 36,068 | 50,430 | 6,799 | 1,539,872 | 189,755 | 4,870 |
| La567 | 3,165,906,177 | 52,962 | 28,528 | 59,777 | 5,431 | 2,136,670 | 259,140 | 3,531 |

**Table 4.** BUSCO scores for the *de novo* elephant assemblies.

| **Assembly** | **Complete** | **Single** | **Duplicated** | **Fragmented** | **Missing** | **Total** |
| --- | --- | --- | --- | --- | --- | --- |
| Em538 (10x+Hi-C) | 8,418 (91.3%) | 8,272 (89.7%) | 146 (1.6%) | 147 (1.6%) | 661 (7.1%) | 9,226 |
| Em538 (10x) | 8,383 (90.9%) | 8,236 (89.3%) | 147 (1.6%) | 168 (1.8%) | 675 (7.3%) | 9,226 |
| Em145 | 8,302 (89.9%) | 8,243 (89.3%) | 59 (0.6%) | 197 (2.1%) | 727 (8.0%) | 9,226 |
| Em146 | 8,346 (90.4%) | 8,243 (89.3%) | 103 (1.1%) | 161 (1.7%) | 719 (7.9%) | 9,226 |
| Em165 | 8,314 (90.1%) | 8,227 (89.2%) | 87 (0.9%) | 192 (2.1%) | 720 (7.8%) | 9,226 |
| Em166 | 8,361 (89.2%) | 8,227 (89.2%) | 134 (1.5%) | 170 (1.8%) | 695 (7.5%) | 9,226 |
| La66 | 8,408 (91.1%) | 8,358 (90.6%) | 50 (0.5%) | 161 (1.7%) | 657 (7.2%) | 9,226 |
| La143 | 8,414 (91.2%) | 8,302 (90.0%) | 112 (1.2%) | 148 (1.6%) | 664 (7.2%) | 9,226 |
| La241 | 8,376 (90.8%) | 8,331 (90.3%) | 45 (0.5%) | 161 (1.7%) | 689 (7.5%) | 9,226 |
| La567 | 8,416 (91.2%) | 8,363 (90.6%) | 53 (0.6%) | 163 (7.0%) | 647 (7.0%) | 9,226 |

### Repeat-masking and genome mappability

We identified repeat elements in the Em538 10x+Hi-C assembly using RepeatMasker v. 4.0.9 (using RMBlast v. 2.9.0+) and RepeatModeler v. 2.0.1 (Flynn et al. 2020; Smit et al. 2013–2015). First, we annotated repeats in the assembly using the Afrotheria repeat database (RepeatMasker options: ‘-gccalc -nolow -xsmall -species afrotheria’). Next, we used the initial repeat-masked genome to build a custom Asian elephant repeat database using RepeatModeler. We then produced a final repeat-masked genome by rerunning RepeatMasker on the Em538 10x+Hi-C assembly using the custom repeat database. We calculated the (30,2)-mappability of the Em538 10x+Hi-C assembly using GenMap v. 1.2.0 with SeqAn v. 2.4.1 (‘genmap map -K30 -E2’: Pockrandt et al. 2020; Reinert et al. 2017).

### Genome annotation

We downloaded all annotated peptide and cDNA sequences for the African savannah elephant reference assembly (loxAfr3.0: Broad Institute 2009, NCBI Accession GCA_000001905.1) from Ensembl (versions dated 26 July 2022: http://useast.ensembl.org/Loxodonta_africana/Info/Annotation). We aligned peptide sequences to the soft-masked Em538 10x+Hi-C assembly using Exonerate v. 2.2.0 (Slater and Birney 2005) (options ‘--model protein2genome --showtargetgff T’). We used BLAT v. 36 (Kent, 2002) to map cDNAs to the masked Asian elephant assembly with a minimum identity of 92% (option ‘-minIdentity=92’). We then generated hints for gene prediction from these mappings using the exonerate2hints.pl (peptides) and blat2hints.pl (cDNAs) scripts from the AUGUSTUS 3.3.2 package (Stanke et al. 2006a, 2006b, 2008). We used these hints, along with the Em538 10x+Hi-C-assembly-specific retraining parameters from the BUSCO analysis, to predict genes on both strands of the repeat-masked Em538 10x+Hi-C assembly using AUGUSTUS (options ‘--softmasking=1 --strand=both --singlestrand=true --extrinsicCfgFile=extrinsic.E.XNT.RM.cfg -- alternatives-from-evidence=true --gff3=on --uniqueGeneId=true’). We functionally characterized the AUGUSTUS-predicted genes using InterProScan v. 5.60-92.0 with all available eukaryotic databases and outputting gene ontology terms (option ‘-goterms’) (Jones et al. 2008).

### Germline mutation rate

We estimated genomic germline *de novo* mutation rates using RatesTools v. 0.5.13 (Armstrong and Campana 2023). The Asian elephant trio samples (sire: Em147; dam: Em165; offspring: Em538) were aligned to the Em538 10x+Hi-C assembly, while the African savannah elephants (sire: La368; dam La241; offspring: La567) were aligned to the loxAfr4.0 reference (Palkopoulou et al. 2018). The Em538 10x+Hi-C assembly has not been anchored to chromosomes, so all sequences longer than 100,000 bp before site- and region-filtration and longer than 10,000 bp after filtration were retained in the analysis. We retained only sequences assigned to autosomes for the African elephant analysis. Sites were filtered to depths (DP) between 20 and 250 and a minimum genotype quality (GQ) of 65. Otherwise, RatesTools pipeline filtering and program versions were as described in Armstrong and Campana (2023) for the wolf (*Canis lupus*) and chimpanzee (*Pan troglodytes*) datasets analyzed therein.

### Genomic population history

We aligned sequences using a custom Nextflow (Di Tommaso et al. 2017) pipeline (https://github.com/campanam/Elephants). We aligned the 10x- and Illumina resequencing reads to the appropriate reference genome (Em538 10x+HiC for Asian elephants and loxAfr4.0 [ftp://ftp.broadinstitute.org/pub/assemblies/mammals/elephant/loxAfr4/] for African savannah elephants: Palkopoulou et al. 2018) using BWA-MEM v. 0.7.17-r1188 (Li 2013). We used SAMtools v. 1.13 (Danecek et al. 2021; Li et al. 2009) to extract mapped reads (‘samtools view -F 4’), fix mate pair tags (‘samtools fixmate -r -m’), and sort the alignments by coordinate (‘samtools sort’). We left-aligned indels for each of the resulting alignments using the Genome Analysis Toolkit (GATK) v. 4.3.0.0 LeftAlignIndels command (McKenna et al. 2010) and marked duplicates using SAMtools (‘samtools markdup’). We calculated alignment statistics using SAMtools (‘samtools flagstat’) and discarded any alignment with fewer than 10 unique aligned reads. Since Em538 was sequenced on two separate lanes, we then merged its libraries using SAMtools (‘samtools merge’) and then re-left-aligned, marked duplicates and calculated alignment statistics for the merged libraries as above. A genome consensus sequence was generated for each elephant using BCFtools v 1.13 (Danecek et al. 2021) and the vcfutils.pl script, requiring a minimum mapping quality of 20, a minimum read depth across all sites of 20 for Em538 and 10 for all other elephants, and a maximum read depth of all sites of 120 for Em538 and 60 for all other elephants (‘bcftools mpileup -q 20 --ignore-RG’, ‘bcftools call -c’ and ‘vcfutils.pl vcf2fq -d <minimum depth> -D <maximum depth>’ commands).

We then reconstructed the demographic history for each elephant using the Pairwise Sequential Markovian Coalescent (PSMC: Li and Durbin 2011). The consensus sequences were converted to psmcfa format using the fq2psmca program from PSMC v. 0.6.5 (Li and Durbin 2010), requiring a minimum quality of 20 (option ‘-q 20’). We performed PSMC on the psmcfa file (options ‘-N25 -t15 -r5 -p “4+25*2+4+6”’). We bootstrapped the PSMC using 100 bootstrap replicates by splitting the original psmcfa file using the splitfa program and then running psmc on the replicates under the same conditions. Results were visualized for each elephant with psmc_plot.pl on the concatenated bootstrap files using a generation time of 25 years (Williams et al. 2020; Wittemyer et al. 2013) and the genomic mutation rates calculated from the RatesTools analyses above (5.3 × 10^-9^ and 3.9 × 10^-9^ substitutions/site/generation for Asian and African elephants respectively) (Figures S1–S2).

### Bait design

To generate a bait set for Asian elephant population genomics from low-quality samples, we identified an initial set of variants that varied both among and between elephant species from our set of nine 10x-sequenced genomes. First, we masked the initial Em538 10x assembly using RepeatMasker v. 4.0.7 (Smit et al. 2013–2015) using a library of known mammalian repeats (option ‘-species mammalia’). We trimmed the 10x sequence data using Trimmomatic v. 0.36 (Bolger et al. 2014) in paired-end mode (options ‘ILLUMINACLIP:TruSeq3-PE.fa:2:30:10 LEADING:3 TRAILING:3 SLIDINGWINDOW:4:15 MINLEN:36’). We then realigned the trimmed 10x sequence data to the Em538 10x assembly using BWA-MEM v. 0.7.17-r1188 (Li 2013). We used SAMtools 1.6 to fix sequencing mate pairs, sort aligned sequences by coordinate and mark PCR duplicates (Danecek et al. 2021; Li et al. 2009). Alignments were indexed using Picard Tools v. 2.5.0 (Broad Institute 2016), and indels were left-aligned using the GATK v. 4.0.8.1 LeftAlignIndels command (McKenna et al. 2010). We joint-genotyped all nine 10x-sequenced elephant individuals together. We called variants using the BCFtools v. 1.6 (Danecek et al. 2021) commands ‘mpileup’ (options ‘-C 50 -d 250 -- ignore-RG’) and ‘call’ (options ‘-m -v’). We retained SNP sites with quality scores ≥ 20 using the BCFtools ‘filter’ command (option ‘-i ‘TYPE=“snp” && QUAL>=20’’). We extracted SNPs that were polymorphic within Asian elephants and removed sites with a minor allele frequency less than 0.05 using VCFtools v. 0.1.14 (Danecek et al. 2011). Finally, we thinned SNPs less than 10,000 bp apart using VCFtools v. 0.1.17. Using BaitsTools (Campana, 2018) v. 1.6.6 (‘vcf2baits -t 200000 -d 20000 -j -O 20 -k 4 -l -B -D -w --disable-lc -c -N -n 30.0 -x 50.0 -K 25.0 -J 4’), we generated 61,551 candidate baits for a custom myBaits kit (Daicel Arbor Biosciences). Following further “stringent” filtration using the myBaits pipeline (Stringent=pass, %RM=0.0, TopHitRM=0.0, HetDimers=0.0, dG ≥ −7.0, BLAST-Hits=1), 38,243 candidate baits covering 27,225 SNP sites (mean tiling depth 1.40×) remained. We selected a final 20,000 bait set using a custom Ruby script elephant_drop_baits.rb. The script optimized tiling density by first randomly removing low-coverage SNPs below a specified target tiling threshold (here 2×) until the maximum requested bait count was reached. If there were still too many baits after removal of low coverage sites, the script downsampled baits for over-represented sites. Here, removal of low-coverage sites was sufficient and no higher-coverage sites were downsampled. The final bait set covered 8,982 unique SNP sites with a mean tiling depth of 2.23× (range 1×-3×). The elephant_drop_baits.rb script and final bait set are available on GitHub (https://github.com/campanam/bait-development/tree/main/Elephant) under the Smithsonian Institution’s Terms of Use (https://www.si.edu/termsofuse).

## Results

### Genome assembly

Basic scaffold and contig sequencing statistics are provided in Tables 2 and 3 respectively. See Supplemental Table 2 for the complete Assemblathon 2 genome assembly statistical output (Bradnam et al. 2013). Genome sizes ranged between 3.23 and 3.29 Gb (Table 2). The 10x-only assemblies had scaffold N50s ranging between 22.8 and 29.4 Mb and contig N50s between 101.9 and 692.0 Kb. For the 10x-only assemblies, the longest scaffolds ranged between 76.4 and 160.2 Mb and the longest contigs ranged in length between 860 Kb and 4.27 Mb. The Em538 10x+HiC assembly had a scaffold N50 of 102.3 Mb and a contig N50 of 681.5 Kb. Its longest scaffold and contig were 228.6 Mb and 4.27 Mb respectively. Gaps comprised 0.9 to 5.1% of the assemblies. All assemblies had high BUSCO scores, with between 89.2% and 91.3% complete BUSCOs found (Table 4).

### Genome mappability and annotation

1,281,630,361 bp (40.0%) of the Em538 10x+Hi-C and 1,411,046,220 bp (44.2%) of the loxAfr4.0 genome had a (30,2)-mappability less than 1.0 (i.e. more than one unique genome hit per sequence). 1,673,109,844 bp (51.8%) of the Em538 10x+Hi-C and 1,613,230,813 bp (49.3%) of the loxAfr4.0 genome were annotated as repeat regions (Supplementary Table 3). Repeats were predominantly LINEs (Em538 10x+Hi-C: 36.2%; loxAfr4.0: 34.0%), SINEs (Em538 10x+Hi-C: 3.4%; loxAfr4.0: 4.0%) and LTRs (Em538 10x+Hi-C: 6.4%; loxAfr4.0: 6.6%). AUGUSTUS predicted 63,327 genes (65,892 predicted transcripts) in the Em538 10x+Hi-C assembly, of which 6,104 of the predicted transcripts were fully supported by the provided hints and another 18,385 transcripts were partially supported. AUGUSTUS gene predictions and InterProScan functional analyses are provided in the Smithsonian Figshare repository (https://dx.doi.org/10.25573/data.22041383).

### Germline mutation rate

RatesTools identified nine Asian candidate *de novo* mutations, yielding a point-estimate germline mutation rate of 7.0 × 10^-9^ substitutions/site/generation (block bootstrap mean: 5.3 × 10^-9^; 95% C.I.: 4.3 – 6.2 × 10^-9^). Among the nine candidates, there were two blocks of SNPs: one on scaffold 1 with two SNPs separated by 3 bp and a second on scaffold 5 where three SNPs were found within 28 bp. These most likely represent either single complex *de novo* mutation events (Chan and Gordenin 2015) or errors due to sequence misalignment. We therefore consider the bootstrapped mean value (5.3 × 10^-9^) as a better estimate of the Asian elephant mutation rate. RatesTools identified four African savannah elephant candidate *de novo* mutations, yielding a point-estimate germline mutation rate of 3.9 × 10^-9^ substitutions/site/generation (block bootstrap mean: 4.0 × 10^-9^; 95% C.I.: 3.2 – 4.8 × 10^-9^).

### Demographic history

Our reconstructed demographic histories of Asian and African savannah elephants largely agreed with those of Palkopoulou et al. (2018). Each had distinct species-specific demographic curves, and the shapes of the curves were very similar to their previously published PSMC results. The primary difference between them was the timing of demographic events, which can be explained by the use of different mutation rates and generation times in the two studies.

## Discussion

In comparison to the Icky assembly by Tollis et al. (2021), our reference Em538 10x+Hi-C assembly represents a 37× improvement in scaffold N50 (102.3 Mb versus 2.77 Mb for Icky) and 8.5× improvement in contig N50 (681.5 Kb versus 79.8 Kb). Our 10x Asian elephant *de novo* genome assemblies are also more contiguous than the current Icky assembly (8.2 – 10.2× scaffold N50; 1.3 – 8.7× contig N50). Furthermore, the Em538 10x+Hi-C assembly has a similar scaffold N50 to the DNA Zoo Hi-C assembly (1.1×: 102.3 Mb versus 96.0 Mb), but a much larger contig N50 (11.8×: 681.5 Kb versus 57.6 Kb). Conversely, the current VGP primary assembly (mEleMax1.pri.cur.20220616) has a similar scaffold N50 (117 Mb: 1.1× Em538 10x+Hi-C), but much larger contig N50 (77 Mb: 113×). Regardless of the variation in genome qualities, the addition of our nine *de novo* elephant assemblies will facilitate research on elephant health, population structure, and evolutionary genomics. This information will also be critical in increasing our appreciation and understanding of the role of genome rearrangements in evolution and disease (which cannot be thoroughly investigated using short-read resequencing) (Schuy et al. 2022).

Germline mutation rates are a critical parameter for reconstructing demographic histories and modeling evolutionary processes (Armstrong and Campana 2023). Previously, based on the fossil-calibrated divergence of *Loxodonta* and *Elephas*, Palkopoulou et al. (2018) estimated a mutation rate for Elephantidae of 0.406 × 10^-9^ substitutions/site/year (uncertainty range: 0.213 × 10^-9^ – 0.599 × 10^-9^ substitutions/site/year). Assuming a generation time of 25 years across the clade, this rate is equivalent to 10.1 × 10^-9^ substitutions/site/generation (range: 5.3 × 10^-9^ – 15.0 × 10^-9^ substitutions/site/generation). While our point estimates of germline mutation rates are slightly slower than the rate used in Palkopoulou et al. (Asian: 5.3 × 10^-9^ substitutions/site/generation; African savannah: 3.9 × 10^-9^ substitutions/site/generation), the trio-based and fossil-calibrated rates show remarkable concordance (less than an order of magnitude with overlap in the estimate confidence intervals) despite the different methodologies, assumptions, and sources of error. Moreover, our estimates are based on single trios for each species, which likely underestimates the range of mutation rates among elephant trios (Bergeron et al. 2022).

Whole genome analyses are impractical in many elephant range countries due to the difficulty in obtaining and processing fresh tissues and/or blood samples (e.g. de Flamingh et al. 2023). Non-invasive samples are much easier to obtain, store and analyze and have been demonstrated to be an important tool in conservation efforts of African and Asian elephants (Ahlering et al. 2020). Our in-solution capture assay presents a cost-effective approach to genotype thousands of genomic markers in a single assay and is particularly well-suited for non-invasive samples (e.g. Campana et al. 2016; Parker et al. 2022). Additionally, because of the flexibility of capture methods, additional markers of interest can easily be incorporated into the bait set presented here, allowing for broad application of this resource to elephant conservation. Our capture array approach offers a standardized method that can be used to accurately guide *ex-situ* and *in-situ* conservation management programs, thereby allowing for comparisons across elephant range countries and breeding programs that can help safeguard genetic diversity in dwindling elephant populations worldwide.

While the resources we present here should allow for more detailed and uniform studies, its ultimate purpose is to be deployed in a large-scale project largely focused on elephant health, such as elephant endotheliotropic herpes virus (EEHV), the leading cause of death in captive Asian elephant calves world-wide (Long et al. 2016; Seilern-Moy et al. 2016; Kendall et al. 2016; Bhusri et al. 2017; Bronson et al. 2017). By combining these genomic resources, with recently published epigenetic clocks and methylation studies in elephants (Prado et al. 2021), and existing datasets on individual elephant temperament (Prado et al. 2020), social life (Prado-Oviedo et al. 2016), EEHV and health histories (Edwards et al. 2019), we aim to identify species-specific genetic variants and genes that could possibly influence disease and morbidity in these species. We hope that the results of these multidisciplinary studies will have significant conservation applications such as a better understanding of the genetic underpinnings of immune system function and disease susceptibility in elephants, as well as act as foundational data for the development of species-specific interventions such as vaccines and antivirals. It will also have a broad impact on how disease is studied and managed for elephants globally, empowering elephant managers, veterinarians and owners to proactively address EEHV and other diseases in a comprehensive manner in order to maximize breeding efforts and optimize health throughout an elephant’s life.

## Acknowledgements

This study was approved by the NZCBI IACUC (# 20-17), and all participating zoos and endorsed by the elephant TAG/SSP. We acknowledge Erin Latimer (NZCBI) and Bethany Nelson (NZCBI) and the following zoos’ staffs for support of the project: Calgary Zoo, Canada; Indianapolis Zoo, National Zoological Park and Smithsonian Conservation Biology Institute, Oklahoma City Zoo, Pittsburgh Zoo and Aquarium, and Utah’s Hogle Zoo. We thank Brian Brunelle (Daicel Arbor Biosciences) for assistance in producing the Asian elephant bait set and Mark Daly (Dovetail Genomics) for providing the Dovetail Hi-C library kit. Computing was performed on the Smithsonian High Performance Computing (SI/HPC) cluster ‘Hydra’, Smithsonian Institution (https://doi.org/10.25572/SIHPC).

## Data Availability

We have deposited the primary data underlying these analyses as follows:

□ DNA sequences: Genbank accessions JAOTIA000000000, JAOTIA020000000, JAOTIB000000000, JAOTIC000000000, JAOTHU000000000, JAOTHV000000000, JAOTHW000000000, JAOTHX000000000, JAOTHY000000000, JAOTHZ000000000; NCBI BioProject PRJNA840935; NCBI BioSamples SAMN28571074–SAMN28571084; NCBI SRA SRR26017082–SRR26017093.
□ Genome annotation: Smithsonian Figshare DOI: 10.25573/data.22041383
□ Custom code: GitHub https://github.com/campanam/Elephants, https://github.com/campanam/genome_decontaminate, https://github.com/campanam/bait-development/tree/main/Elephant
□ Bait sequences: GitHub https://github.com/campanam/bait-development/blob/main/Elephant/elephant_baits.fa

## Supplementary Material

**Supplementary Table 1.**
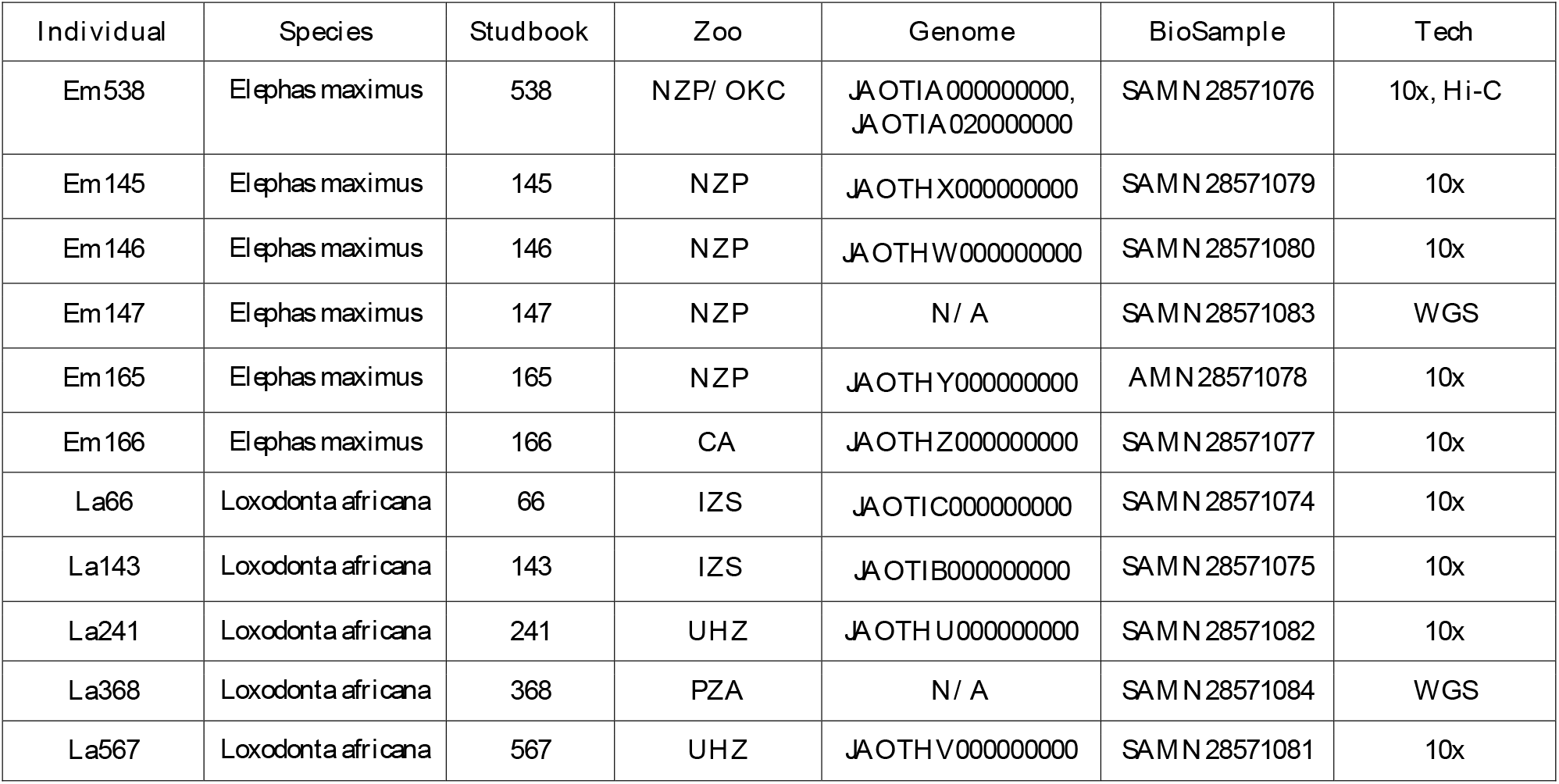
Elephant sample metadata. The ‘Studbook’ column lists the AZA Elephant studbook number. The ‘Zoo’ column lists the home zoo(s) of the individual (‘CZ’ = Calgary Zoo, Calgary, AB, Canada; ‘IZS’ = Indianapolis Zoological Society, Indianapolis, IN; ‘NZP’ = Smithsonian’s National Zoo and Conservation Biology Institute, Washington, DC; ‘OKC’ = Oklahoma City Zoo, Oklahoma City, OK; ‘PZA’ = Pittsburgh Zoo & Aquarium, Pittsburgh, PA; ‘UHZ’ = Utah’s Hogle Zoo, Salt Lake City, UT). The ‘Genome’ and ‘BioSample’ columns list NCBI accessions for the genome assemblies and BioSamples. The ‘Tech’ column lists sequencing technologies (‘10x’ = 10x Chromium; ‘Hi-C’ = Dovetail Hi-C; ‘WGS’ = whole genome shotgun resequenced).

**Supplementary Table 2.** Complete statistics for the *de novo* elephant genome assemblies as assessed using the assemblathon_stats.pl script from the Assemblathon 2 challenge (Bradnam et al. 2013).

**Supplementary Table 3:** Fraction of the Em538 10x+HiC and loxAfr4.0 genomes comprised of repetitive elements as annotated by RepeatMasker.

| Repeat Class | Em538 10x+HiC | loxAfr4.0 |
| --- | --- | --- |
| SINEs | 108,449,705 bp (3.36%) | 132,069,390 bp (4.04%) |
| LINEs | 1,170,806,702 bp (36.24%) | 1,113,466,007 bp (34.03%) |
| LTRs | 205,802,719 bp (6.37%) | 214,792,061 bp (6.56%) |
| DNA elements | 89,463,042 bp (2.77%) | 68,147,076 bp (2.08%) |
| Unclassified repeats | 69,179,223 bp (2.14%) | 75,771,394 bp (2.32%) |
| Small RNAs | 15,369,900 bp (0.48%) | 1,371,161 bp (0.04%) |
| Satellites | 35,443,555 bp (1.10%) | 27,126,252 bp (0.83%) |
| Simple repeats | 1,441,332 bp (0.04%) | 1,196,056 bp (0.04%) |

**Supplementary Figure 1**: Demographic history reconstructions for Asian elephants.

**Supplementary Figure 2**: Demographic history reconstructions for African savannah elephants.

## References

Ahlering M, Budd K, Schuttler S, Eggert LS. 2020. Genetic analyses of non-invasively collected samples aids in the conservation of elephants. In: Ortega J, Maldonado JE(eds.). Conservation Genetics in Mammals: Integrating Research using Novel Approaches. Springer Cham, Pp. 229–248.

Ahlering MA, Eggert LS, Western D, Estes A, Munishi L, Fleischer R, Roberts M, Maldonado JE. 2012a. Identifying source populations and genetic structure for Savannah Elephants in human-dominated landscapes and protected areas in the Kenya-Tanzania borderlands. PLoS One. 7(12):e52288.

Ahlering MA, Maldonado JE, Fleischer, RC, Western D, Eggert LS. 2012b. Fine-scale group structure and demography of African Savanna Elephants recolonizing lands outside protected areas. Divers Distri. 18:952–996.

Aiden Lab. 2022. Asian elephant (Elephas maximus) [accessed 2023 Jan 27]. Available from: https://www.dnazoo.org/assemblies/Elephas_maximus

Armstrong EE, Campana MG. 2023. RatesTools: a Nextflow pipeline for detecting *de novo* germline mutations in pedigree sequence data. Bioinformatics. 39(1):btac784.

Athira TK, Vidya TNC. 2021. Elephant social systems: what do we know and how have molecular tools helped? J Indian Inst Sci. 101:257–278.

Bergeron LA, Besenbacher S, Turner T, Versoza CJ, Wang RJ, Price, AL, Armstrong E, Riera M, Carlson J, Chen H-y, et al. 2022. The Mutationathon highlights the importance of reaching standardization in estimates of pedigree-based germline mutation rates. eLife. 11:e73577.

Bhusri B, Suksai P, Mongkolphan C, Tiyanun E, Tatanakor P, Chaichoun K, Sariya L. 2017. Detection of elephant endotheliotropic herpesvirus 4 in captive Asian elephants (*Elephas maximus*) in Thailand. Thai J Vet Med. 47:97–102.

Blake S, Hedges S. 2004. Sinking the flagship: the case of forest elephants in Asia and Africa. Conserv Biol. 18(5):1191–1202.

Bolger AM, Lohse M, Usadel B. 2014. Trimmomatic: a flexible trimmer for Illumina sequence data. Bioinformatics. 30(15):2114–2120.

Bradnam KR, Fass JN, Alexandrov A, Baranay P, Bechner M, Birol I, Boisvert S, Chapman JA, Chapuis G, Chikhi R, et al. 2013. Assemblathon 2: evaluating *de novo* methods of genome assembly in three vertebrate species. Gigascience. 2:10.

Broad Institute. 2009. Elephant genome project. [accessed 2023 January 24]. Available from: https://www.broadinstitute.org/elephant/elephant-genome-project

Broad Institute. 2016. Picard version 2.5.0. [accessed 2018 August 8]. Available from: https://broadinstitute.github.io/picard

Bronson E, McClure M, Sohl J, Wiedner E, Cox S, Latimer EM, Pearson VR, Hayward GS, Fuery A, Ling PD. 2017. Epidemiologic evaluation of elephant endotheliotropic herpesvirus 3b infection in an African elephant *(Loxodonta africana)*. J. Zoo Wild. Med. 48: 335–343.

Brown JL. 2014. Comparative reproductive biology of elephants. In: Holt WV, Brown, JL, Comizolli P (eds.). Reproductive Sciences in Animal Conservation: Progress and Prospects. Advances in Experimental Medicine and Biology, No. 753. New York, NY: Springer Science and Business Media, Pp. 135–169.

Campana MG. 2018. BaitsTools: Software for Hybridization Capture Bait Design. Mol Ecol Resour. 18(2):356–61.

Campana MG. 2019. genome_decontaminate. [accessed 2022 August 18]. Available from: https://github.com/campanam/genome_decontaminate

Campana MG, Hawkins MTR, Henson LH, Stewardson K, Young H S, Card LR, Lock J, Agwanda B, Brinkerhoff J, Gaff HD, et al. 2016. Simultaneous identification of host, ectoparasite and pathogen DNA via in-solution capture. Mol Ecol Resour. 16(5):1224–1239.

Chan K, Gordenin DA. 2015. Clusters of multiple mutations: incidence and molecular mechanisms. Annu Rev Genet. 49:243–267.

Chase MJ, Schlossberg S, Griffin CR, Bouche PJC, Djene SW, Elkan PW, Ferreira S, Grossman F, Kohi EM, Landen K, et al. 2016. Continent-wide survey reveals massive decline in African savannah elephants. PeerJ 4:e2354.

Clubb R, Mason G. 2002. A Review of the Welfare of Zoo Elephants in Europe: A report commissioned by the RSPCA. Oxford: University of Oxford, Animal Behaviour Research Group. Convention on.International Trade in Endangered Species of Wild Fauna and Flora (CITES). 2023. Appendices I, II, and III valid from 11 January 2023 [accessed 2023 January 27]. Available from: http://www.cites.org/eng/app/appendices.php

Danecek P, Auton A, Abecasis G, Albers CA, Banks E, DePristo MA, Handsaker RE, Lunter G, Marth GT, Sherry ST, et al.; 1000 Genomes Project Analysis Group. 2011. The variant call format and VCFtools. Bioinformatics. 27:2156–2158.

Danecek P, Bonfield JK, Liddle J, Marshall J, Ohan V, Pollard MO, Whitwham A, Keane T, McCarthy SA, Davies RM, et al. 2021. Twelve Years of SAMtools and BCFtools. GigaScience 10(2):giab008.

de Flamingh A, Ishida Y, Pečnerová P, Vilchis S, Siegismund HR, van Aarde RJ, Malhi RS, Roca AL. 2023. Combining methods for non-invasive fecal DNA enables whole genome and metagenomic analyses in wildlife biology. Front Genet.13:1021004.

Di Tommaso P, Chatzou M, Floden EW, Prieto Barja P, Palumbo E, Notredame C. 2017. Nextflow enables reproducible computational workflows. Nat Biotechnol. 35:316–319.

Edwards KL, Miller MA, Carlstead K, Brown JL. 2019. Relationships between housing and management factors and clinical health events in elephants in North American zoos. PLoS ONE 14(6), e0217774.

Faust LJ, Marti K. 2011a. Technical report on zoo risk modeling of the North American African elephant SSP population. Chicago: Lincoln Park Zoo.

Faust LJ, Marti K. 2011b. Technical report on zoo risk modeling of the North American Asian elephant SSP population. Chicago: Lincoln Park Zoo.

Fleischer RC, Perry EA, Muralidharan K, Stevens EE, Wemmer CM. 2001. Phylogeography of the Asian elephant *(Elephas maximus)* based on mitochondrial DNA. Evolution. 55:1882–1892.

Flynn JM, Hubley R, Goubert C, Rosen J, Clark AG, Feschotte C, Smit AF. 2020. RepeatModeler2 for automated genomic discovery of transposable element families. Proc Natl Acad Sci U S A. 117(17):9451–9457.

Gobush KS, Edwards CTT, Balfour D, Wittemyer G, Maisels F, Taylor RD. 2022. *Loxodonta africana* (amended version of 2021 assessment). The IUCN Red List of Threatened Species. 2022:e.T181008073A223031019.

Gobush KS, Edwards CTT, Maisels F, Wittemyer G, Balfour D, Taylor RD. 2021. *Loxodonta cyclotis* (errata version published in 2021). The IUCN Red List of Threatened Species. 2021:e.T181007989A204404464.

Gobush KS, Mutayoba BM, Wasser SK. 2008. Long-term impacts of poaching on relatedness, stress physiology, and reproductive output of adult female African elephants. Conserv Biol. 22:1590–1599.

Hauf J, Waddell PJ, Chalwatzis N, Joger U, Zimmermann FK. 2000. The complete mitochondrial genome sequence of the African elephant *(Loxodonta africana)*, phylogenetic relationships of Proboscidea to other mammals and D-loop heteroplasmy. Zoology. 102:184–195.

Helmy M, Awad M, Mosa KA. 2016. Limited resources of genome sequencing in developing countries: challenges and solutions. Appl Trans Genom. 9:15–19.

Hildebrandt TB, Hermes R, Saragusty J, Potier R, Schwammer HM, Balfanz F, Vielgrader H, Baker B, Bartels P, Göritz F. 2012. Enriching the captive elephant population genetic pool through artificial insemination with frozen-thaw semen collected in the wild. Theriogenology. 78:1398–1404.

Jones P, Binns D, Chang HY, Fraser M, Li W, McAnulla C, McWilliam H, Maslen J, Mitchell A, Nuka G, et al. 2014. InterProScan 5: genome-scale protein function classification. Bioinformatics. 30:1236–1240.

Kendall R, Howard L, Masters N, Grant, R. 2016. The impact of elephant endotheliotropic herpesvirus on the captive Asian elephant (Elephas maximus) population of the United Kingdom and Ireland (1995-2013). J Zoo Wild Med. 47:1–14.

Kent WJ. 2002. BLAT–the BLAST-like alignment tool. Genome Res. 12:656–664.

Lei R, Brenneman RA, Louis, Jr. EE. 2008. Genetic diversity in the North American captive African elephant collection. J Zoo. 275:252–267.

Lei R, Brenneman RA, Schmitt DL, Louis, Jr. EE. 2011. Genetic diversity in the North American captive Asian elephants. J Zoo. 286:38–47.

Leimgruber P, Oo ZM, Aung M, Kelly DS, Wemmer C., Senior B, Songer M. 2011. Current status of elephants in Myanmar. Gajah. 35:76–86.

Leimgruber P, Gagnon JB, Wemmer C, Kelly DS, Songer MA, Selig ER. 2003. Fragmentation of Asia’s remaining wildlands: implications for Asian elephant conservation. Anim Conserv. 6(4):347–359.

Li H. 2013. Aligning sequence reads, clone sequences and assembly contigs with BWA-MEM. arXiv. 1303.3997.

Li H, Durbin R. 2011. Inference of human population history from individual whole-genome sequences. Nature. 475:493–496.

Li H, Handsaker B, Wysoker A, Fennell T, Ruan J, Homer N, Marth G, Abecasis G, Durbin R: 1000 Genome Project Data Processing Subgroup. 2009. The sequence alignment/map format and SAMtools. Bioinformatics. 25:2078–2079.

Long, S.Y., Latimer, E.M., Hayward, G.S. 2016. Review of elephant endotheliotropic herpesviruses and acute hemorrhagic disease. ILAR J. 56, 283–296.

Maisels F, Strindberg S, Blake S, Wittemyer G, Hart J, Williamson E., Aba’a R, Abitsi G, Ambahe RD, Amsini F, et al. 2013. Devastating decline of forest elephants in Central Africa. PLoS One. 8(3):e59469.

Manni M, Berkeley MR, Seppey M, Simão FA, Zdobnov EM. 2021. BUSCO update: novel and streamlined workflows along with broader and deeper phylogenetic coverage for scoring of Eukaryotic, Prokaryotic, and viral genomes. Mol Biol Evol.38(10):4647–4654.

McKenna A, Hanna M, Banks E, Sivachenko A, Cibulskis K, Kernytsky A, Garimella K, Altshuler D, Gabriel S, Daly M, et al. 2010. The genome analysis toolkit: a mapreduce framework for analyzing next-generation DNA sequencing data. Genome Res. 20:1297–1303.

National Center for Biotechnology Information (NCBI). 2022. NCBI Foreign Contamination Screen. [accessed 2023 January 25]. Available from: https://github.com/ncbi/fcs

Olson D, Wiese RJ. 2000. State of the North American African elephant population and projections for the future. Zoo Biol. 9:311–320.

Palkopoulou E, Lipson M, Mallick S, Nielsen S, Rohland N, Baleka S, Karpinski E, Ivancevic AM, To T-H, Kortschak RD, et al. 2018. A comprehensive genomic history of extinct and living elephants. Proc Natl Acad Sci U S A. 115(11):E2566–E2574.

Parker LD, Campana MG, Quinta JD, Cypher B, Rivera I, Fleischer RC, Ralls K, Wilbert TR, Boarman R, Boarman WI, Maldonado JE. 2022. An efficient method for simultaneous species, individual, and sex identification via in-solution single nucleotide polymorphism capture from low-quality scat samples. Mol Ecol Resour. 22(4):1345–1361.

Pockrandt C, Alzamel M, Iliopoulos CS, Reinert K. 2020. GenMap: ultra-fast computation of genome mappability. Bioinformatics. 36:3687–3692.

Prado-Oviedo NA, Bonaparte-Seller MK, Malloy EJ, Meehan CL, Mench JA, Carlstead K, Brown JL. 2016. Evaluations of demographics and social life events of Asian *(Elephas maximus)* and African elephants *(Loxodonta africana)* in North American zoos. PLoS One. 11:e0154750.

Prado NA, Malloy E, Carlstead K, Wielebnowski N, Brown JL. 2020. Ovarian cyclicity and prolactin status of African elephants *(Loxodonta africana)* in North American zoos may be influenced by life experience and individual temperament. Horm. Behav. 125:104804.

Prado NA, Brown JL, Zoller JA, Haghani A, Yao M, Bagryanova LR, Campana MG, Maldonado JE, Raj K, Schmitt D, et al. 2021. Epigenetic clock and methylation studies in elephants. Aging Cell. 20(7):e13414.

Reinert K, Dadi TH, Ehrhardt M, Hauswedell H, Mehringer S, Rahn R, Kim J, Pockrandt C, Winkler J, Siragusa E, et al. (2017) The SeqAn C++ template library for efficient sequence analysis: A resource for programmers. J Biotechnol.261:157–168.

Roca AL, Georgiadis N, Pecon-Slattery J, O’Brien SJ. 2001. Genetic evidence for two species of elephant in Africa. Science. 293: 1473–1477.

Rogaev EI, Moliaka YK, Malyarchuk BA, Kondrashov FA, Derenko MV, Chumakov I, Grigorenko AP. 2006. Complete mitochondrial genome and phylogeny of a Pleistocene mammoth *Mammuthus primigenius*. PLoS Biol. 4(3):e73.

Rohland N, Reich D, Mallick S, Meyer M, Green RE, Georgiadis NJ, Roca AL, Hofreiter M. 2010. Genomic DNA sequences from mastodon and woolly mammoth reveal deep speciation of forest and savanna elephants. PLoS Biol.8(12):e1000564.

Schuy J, Grochowski CM, Carvalho CMB, Lindstrand A. 2022. Complex genomic rearrangements: an underestimated cause of rare diseases. Trends Genet.38(11):1134–1146.

Seilern-Moy K, Darpel K, Steinbach F, Dastjerdi A. 2016. Distribution and load of elephant endotheliotropic herpesviruses in tissues from associated fatalities in Asian elephants. Virus Res. 220:91–96.

Slater GS, Birney E. 2005. Automated generation of heuristics for biological sequence comparison. BMC Bioinformatics. 6:31.

Smit AFA, Hubley R, Green P. 2013–2015. RepeatMasker open-4.0. [accessed 2017 October 13]. Available from: http://www.repeatmasker.org

Stanke M, Diekhans M, Baertsch R, Haussler D. 2008. Using native and syntenically mapped cDNA alignments to improve de novo gene finding. Bioinformatics.24:637–644.

Stanke M, Schöffmann O, Morgenstern B, Waack S. 2006a. Gene prediction in eukaryotes with a generalized hidden Markov model that uses hints from external sources. BMC Bioinformatics. 7:62.

Stanke M, Tzvetkova A, Morgenstern B. 2006b. AUGUSTUS at EGASP: using EST, protein and genomic alignments for improved gene prediction in the human genome. Genome Biol. 7(Suppl 1):S11.1–S11.8.

Thitaram C. 2012. Breeding management of captive Asian elephant *(Elephas maximus)* in range countries and zoos. Jpn J Zoo Wildl Med. 17:91–96.

Thouless CR, Dublin HT, Blanc JJ, Skinner DP, Daniel TE, Taylor RD, Maisels F, Frederick HL, Bouche P. 2016. African Elephant Status Report 2016: an update from the African Elephant Database. Occasional Paper Series of the IUCN Species Survival Commission, No. 60. IUCN, Gland, Switzerland.

Tollis M, Ferris E, Campbell MS, Harris VK, Rupp SM, Harrison TM, Kiso WK, Schmitt DL, Garner MM, Aktipis CA, et al. 2021. Elephant genomes reveal accelerated evolution in mechanisms underlying disease defenses. Mol Biol Evol. 38(9):3606–3620.

Vertebrate Genomes Project. 2022. Elephas maximus Asiatic elephant. GenomeArk. [accessed 2023 Jan 27]. Available from: https://genomeark.github.io/vgp-all/Elephas_maximus/

Vidya TNC, Varma S, Dang NX, Van Thanh T, Sukumar R. 2007. Minimum population size, genetic diversity, and social structure of the Asian elephant in Cat Tien National Park and its adjoining areas, Vietnam, based on molecular genetic analyses. Conserv Genet. 8:1471–1478.

Weisenfeld NI, Kumar V, Shah P, Church DM, Jaffe DB. 2017. Direct determination of diploid genome sequences. Genome Res. 27(5):757–767.

Wiese RJ. 2000. Asian elephants are not self-sustaining in North America. Zoo Biol.19:299–300.

Williams C, Tiwari SK, Goswami VR, de Silva S, Kumar A, Baskaran N, Yoganand K, Menon V. 2020. Elephas maximus. The IUCN Red List of Threatened Species.2020:e.T7140A45818198.

Wittemyer G, Deballen D, Douglas-Hamilton I. 2013. Comparative demography of an at-risk African elephant population. PloS One. 8(1):e53726.

Wittemyer G, Okello JBA, Rasmussen HB, Arctander P, Nyakaana S, Douglas-Hamilton I, Siegismund HR. 2009. Where sociality and relatedness diverge: the genetic basis for hierarchical social organization in African elephants. Proc Biol Sci.276(1672): 3513–3521.

Zhao C, McCarthy SA, Durbin R. 2023. YaHS: yet another Hi-C scaffolding tool. Bioinformatics. 39(1):btac808.

